# The Effect of Evaluating Self’s Emotions on Frontal Alpha Asymmetry

**DOI:** 10.1101/2023.04.02.535188

**Authors:** Masato Ito, Toru Takahashi, Yuto Kurihara, Rieko Osu

**Author notes:** Corresponding Author: Rieko Osu, PhD., Faculty of Human Sciences, Waseda University, 2-579-15 Mikajima, Tokorozawa Saitama, Japan. Abbreviations: Electroencephalography (EEG), Event-Related Potential (ERP), Event-Related Spectral Perturbations (ERSPs), Frontal Alpha Asymmetry (FAA), International Affective Picture System (IAPS), Self-Assessment Manikin (SAM).

## Abstract

In this research to assess emotions from biometric signals, participants are asked to evaluate the emotions they subjectively experienced in order to confirm whether the assumed emotions were actually elicited. However, the evaluation of emotions is not routinely performed in daily life, and it is possible that this evaluation may alter biological signals. In fMRI studies, evaluation has been shown to activate the amygdala, which is said to be related to emotional expression. However, electroencephalography (EEG) studies do not take into consideration the effects of such evaluations, and it is unclear how these evaluations affect emotion-related brain activity observed in EEG. We hypothesized that emotion evaluations would amplify emotions and c alter Frontal Alpha Asymmetry (FAA), which has been shown to be related to emotional pleasantness and unpleasantness. We suspect this is because in order to evaluate one’s emotions, one must pay attention to one’s internal state, and this self-focused attention has been found to enhance the subjective emotional experience. We measured a 29-channel EEG when presented with unpleasant and highly arousing images from the International Affective Picture System (IAPS) from 40 healthy male and female participants. The results revealed that FAA was significantly lower in the condition in which participants rated their own emotions compared to the condition in which they did not. Similar to fMRI studies, this result indicates that emotion-related brain activity is amplified on an EEG. This paper provides a cautionary note regarding the use of such evaluations in EEG emotion estimation studies.

## 1. Introduction

Various studies have attempted to quantify visible human emotions using objective numerical values. Studies on human emotions were performed using questionnaires in which respondents’ answers were based on categories of emotions (joy, sadness, anger, happiness) or dimensions of emotions (pleasant-unpleasant, aroused-sedated) using a Likert scale. However, in recent years more objective measurement methods using biological signals have been described (Shu et al., 2018). Electroencephalogram (EEG) has been used to estimate emotion, and this approach has the advantage of allowing the prompt and objective observation of emotion providing more information than peripheral signals (Alarcao & Fonseca, 2019; Suhaimi et al., 2020). However, a considerable challenge of validity exists in the application of these techniques, as it has been pointed out that differences may exist between the experimental environment in which the data used to train the models are measured and the everyday environment to which the models are applied (Azari et al., 2020).

In the application of the emotion recognition technology to everyday environments, the act of evaluating emotions is a necessary process in improving the validity and accuracy of recognition models. Evaluation, however, has been shown to alter the subjective emotional experience and emotion-related brain activity in a top-down manner. For example, it has been confirmed that the act of reappraisal, a top-down controlled evaluation of a stimulus, can attenuate the emotional experience of the stimulus (Lazarus & Alfert, 1964; Ochsner & Gross, 2005). This reappraisal has been applied clinically to ameliorate phobias. At the same time, it has also been observed that this reappraisal alters EEG indicators related to emotions (Hajcak & Nieuwenhuis, 2006). Thus, it can be seen that evaluation is an important factor that determines emotional experience and may also alter emotional processing itself.

In recent years, the effects of evaluation itself on brain activity have also been examined. In studies that investigate the mechanism of “attention”, one of the methods used to induce attention to an emotional stimulus is to require the participants to evaluate the stimulus (Chun et al., 2011; Keller, 2011). A comparison of trials with and without evaluation for olfactory pleasant/neutral/unpleasant stimuli revealed that evaluation for the emotion that the stimulus possesses modulates attention to the stimulus, which manifests as amplification of event-related potential (ERP) P300 on EEG (Singh et al., 2019). In addition, fMRI studies of emotion have confirmed that emotion evaluation amplifies activity of emotion-related brain areas such as the amygdala (Lee & Siegle, 2012). However, the effect of evaluation on self-emotion on the EEG in emotion estimation studies remains unclear.

In general, the temporal resolution of EEG is higher than that of fMRI and allows for the identification of very narrow temporal changes (Hennig et al., 2003). In the processing of emotions, evaluation is the process occurring after stimuli are perceived, and it is unclear whether the effect is added in the 1000 or 2000 ms time window used for emotion estimation by EEG. In addition, the temporal resolution of fMRI does not allow us to know whether low-order information processing to stimuli is affected or whether higher-order processing with a time delay of several hundred ms, such as ruminations on stimuli caused by the evaluation process, is affected. Therefore, in this study, we examined whether the condition of emotional subjective evaluation modulates emotion-related EEG immediately after the presentation of emotional stimuli.

This study also seeks to provide neuroscientific evidence of self-focused attention, which directs attention to one’s own emotions and internal changes. Muta & Koshikawa (2015) reported that pre-mood measurement may influence the reaction time of a subsequent emotion word discrimination task. They discussed that prior mood measurement elicited self-focused attention and delayed the response to the stimulus as the reason for this result. Although this self-focused attention has been shown to boost emotion subjectively (Scheier & Carver, 1977), from a neuroscientific perspective, the underlying brain activity is not clearly understood. Therefore, we also examined whether the emotional evaluation has an after-effect on the EEG, which means a lasting effect of evaluation even after quitting it.

Frontal Alpha Asymmetry (FAA) is used as an index of EEG in this study. There are a variety of EEG indexes, and many reports have been published on the relationship between EEG and emotion (Coan & Allen, 2004, Hajcak et al., 2010, Liberati et al., 2015). Of these, FAA is an important index often used in basic and applied research as an indicator representing an aspect of emotion (Davidson, 2004). FAA is grounded in the fact that right frontal activity is associated with avoidance behavior and negative emotions while left frontal activity is associated with approach behavior and positive emotions. FAA, calculated by subtracting left frontal alpha power from right frontal alpha power, is an index that is positively correlated with positive emotions. This indicator has shown similar trends with the results of the fMRI studies (Davidson et al., 1999, Canli et al., 1998). In addition, this indicator has been found to be reproducible in studies from numerous fields (Allen et al, 2018, Reznik & Allen, 2018, Meyer et al, 2015, Smith et al, 2017). Therefore, FAA is presumed to be a valid indicator of pleasant and unpleasant emotions. It is also an index that is often used as an indicator in emotion recognition research, and if the effect of evaluations on FAA is occurring, it could potentially shake up the results of previous studies. Therefore, examining the effects of evaluation on FAA is important both as evidence supporting the validity of previous emotion recognition research. At the same time, it is also important in the sense that it demonstrates the need to consider the effects of evaluation for future estimation of more sophisticated internal states, such as estimation of not only responses to stimuli, but also attitudes toward stimuli and prior context.

Therefore, the present study examined whether subjective evaluation of emotion during emotion elicitation affects FAA and whether the same attitude toward the stimulus as during evaluation persists even after emotion evaluation has ceased and influences the FAA. Two experimental conditions were set for the task: without subjective evaluation (Without Evaluation) condition and with subjective evaluation (With Evaluation) condition. The experimental group participants first conducted the With Evaluation condition, followed by Without Evaluation condition. The control group participants conducted Without Evaluation condition twice. We hypothesized that the experimental group would induce a greater modulation of the FAA than the control group because the participants would experience their emotions more intensely by paying attention to their own emotions and that this effect would persist even after emotion evaluation would cease. We also confirmed the amplitude of P300 to see if the ERP modulation is same as in Singh et al. (2019).

## 2. Materials and Methods

### 2.1. Participants

Forty-one students with no history of psychiatric or neurological disorders participated in the experiment. One participant was excluded because an issue with the program occurred during the experiment and normal responses could not be measured. Therefore, 40 individuals participated in the study (age 21.4 ± 2.7 years (mean ± SD), 17 female). Although the sample size was not determined using statistical methods, the number of participants was set to be comparable to sample sizes used in previous studies (N = 28, 14 control) (Hutcherson et al., 2005) taking into account that some data would be rejected during analysis.. Participants were recruited at the Waseda University, Japan. The following exclusion criteria applied: 1) impairment of any of the senses (hearing, vision, smell, taste and touch) or of hand tasks (i.e. vision could be corrected), 2) who agreed to undergo tests to measure physiological signals, 3) any medication taken within 24 hours, 4) caffeine or alcohol consumption within 12 hours, 5) currently experiencing auditory or visual abnormalities that interfere with daily life, 6) significant sleep deprivation or fatigue within 12 hours, 7) a traumatic event experienced in the previous month combined with feelings of distress triggered by recalling that event, and 8) current participation in another intervention study in any field. These conditions were set to control for factors that may affect the EEG measurements and results of the study and to reduce the burden on the participants as much as possible. Finally, none of the recruited participants declined to participate on grounds of the exclusion criteria set out above.

This experiment was approved by the Waseda University Ethics Review Committee on Research Involving Human Participants and conformed to the Code of Ethics of the World Medical Association (Declaration of Helsinki). All participants signed a consent form to participate in the study, had the right to withdraw from the experiment at any time, and received monetary compensation of 1,500 yen per hour for their participation. In order to avoid informing the participants that the purpose of the experiment was related to emotions, we only explained in advance that the purpose of the experiment was to measure biological signals when presented with visual stimuli, and we avoided the use of the keyword “emotion” in the recruitment stage and during the experiment as much as possible. After the experiment, an explanation of the intent was given.

### 2.2. Visual Stimuli and Presentation

The international affective picture systems (IAPS; Lang et al., 2008) were used for the visual stimuli. A total of 134 images were extracted based on affective valence and arousal levels that were pre-labeled by Lang et al. (2008). The extracted images had low emotional valence (high unpleasant) and high arousal (Figure 1). This refers to images with an arousal level of 5 or higher and an emotional valence of 3.55 or lower based on the distribution of emotional valence and arousal level. Images that were ethically problematic such as erotic or grotesque were visually eliminated. The threshold value of 3.55 for emotional value corresponds to the cut-off for the top 25% points, and the threshold value of 5 for arousal level is the value determined by considering the V-shaped distribution characteristic of IAPS images for the top 25% points. Since wearing the electroencephalograph for a long period of time is burdensome for the participants, the experiment was limited to negative high arousal, which is more likely to produce a response, to shorten the experimental time. From this group of 134 images, 100 images were presented pseudo-randomly in the same order among participants (arousal level = 5.96 ± 0.50, valence level = 2.71 ± 0.59). In the experiment, 100 images were presented in four blocks. The IAPS image groups were divided into Blocks 1 through 4 so as to avoid significant differences in emotional valence or arousal. A one-factor ANOVA analysis between Blocks (Blocks 1, 2, 3, and 4) was conducted for emotional valence and arousal, respectively, and the test results showed that no significant differences were detected in the ratings of the image groups (valence: f (3,96) =0.56, p=0.64, arousal: f (3,96) =0.28, p=0.84).

**Figure. 1.**
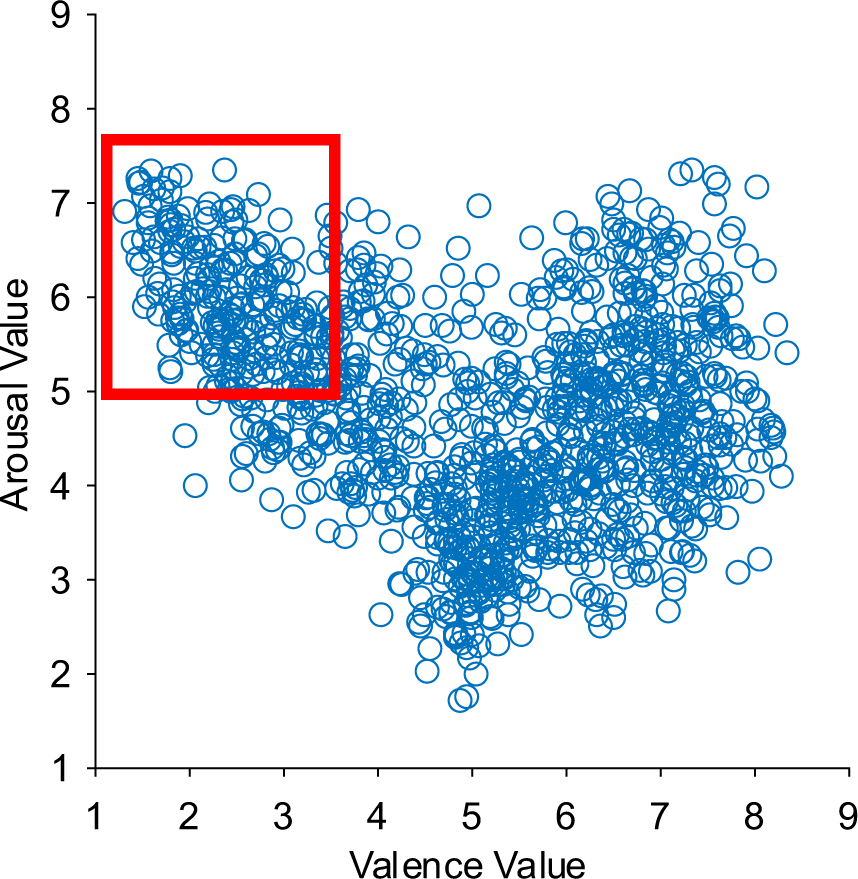
The set of pictures we used for our experiments based on the scatterplot of the label values originally attached to the IAPS pictures. This figure plots the labeling values originally attached to the IAPS images with valence on the horizontal axis and arousal on the vertical axis. Images within the red box (valence<=3.55, arousal>=5) were designated as negative high-arousal images, of which 100 were randomly selected for the experiment.

Visual stimuli were presented using MATLAB’s psychtoolbox (Brainard, 1997; Pelli, 1997; Kleiner et al., 2007) on a 13.3-inch display with a resolution of 2560 x 1600 on a MacBook Pro. The display was positioned approximately 60 cm away from the participant. Participants wore earplugs to control auditory information and sat in a relaxed position in a chair.

### 2.3. Experimental Groups and Conditions

Participants were assigned to two groups, the experimental group, and the control group, in the order of their participation. Because the protocol of the control group required the response data of the experimental group, the assignment was not done randomly, but in the order of first-come-first-served, from the experimental group to the control group.

The experiment was conducted under two conditions: a “With Evaluation” condition in which participants were asked to make a subjective evaluation of their own emotional state (both arousal and valence) after the presentation of each image and a “Without Evaluation” condition in which participants were not asked to make a subjective evaluation of their emotional state (Figure 2A). The experiment was conducted for a total of 4 Blocks (Figure 2B). First, a “Without Evaluation” session was conducted in both groups to fully habituate the participants to the stimuli (Block 1). Then, the experimental group completed the trial under “With Evaluation” conditions and the control group completed the trial under “Without Evaluation” conditions (Block 2). Both groups then participated in a trail under “Without Evaluation” conditions (Block 3). Since the control group only underwent the trial under “Without Evaluation” conditions and there was no reference evaluation value in order to confirm whether the expected emotion was actually elicited or not, a trial under “With Evaluation” conditions was conducted in both groups at the end to collect the evaluation values (Block 4).

**Figure. 2.**
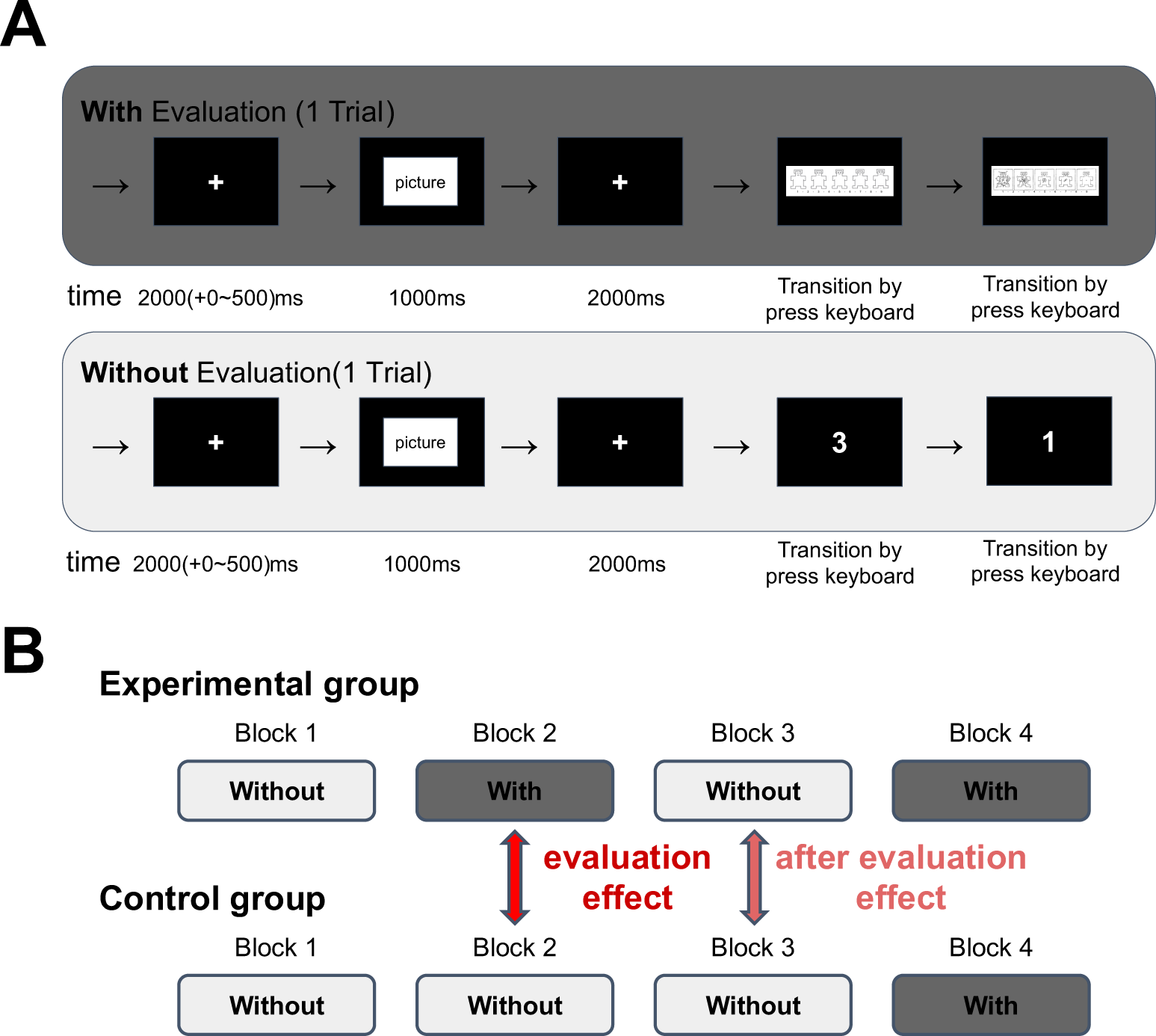
Experimental design. **(A)** The duration of each event for the “With Evaluation” (top) and “Without Evaluation” (bottom) conditions. The white cross was always presented except during visual stimulus presentation and evaluation. The stimulus duration was 1000 ms. The onset of the image stimulus was randomized and appeared 2000 to 2500 ms after the end of the previous trial. Epochs for calculating EEG indexes were extracted from the interval from stimulus to evaluation (−500 ms to 3000 ms after stimulus onset). **(B)** The order of experimental conditions in the experimental and control groups. After a practice trial for both groups under the “Without Evaluation” conditions (Block 1), the experimental group performed the trial under “With Evaluation” conditions and the control group performed the trial under “Without Evaluation” conditions (Block 2). Both groups then performed the trial under “Without Evaluation” condition (Block 3), followed by “With Evaluation” condition (Block 4); the second condition was compared across the groups (control group: “Without Evaluation”, experimental group: “With Evaluation”) to examine the effects of the evaluations, and the third condition was compared across the groups (control group: “Without Evaluation”, experimental group: “Without Evaluation”) to examine the after-effects of the evaluations.

### 2.4. Procedure

The “With Evaluation” condition involved an initial fixation phase in which participants focused on a white cross (randomized from 2000 to 2500 ms based on Matlab’s rand function) (Figure 2A). Next, the stimulus phase consisted of a 1000 ms presentation of the IAPS emotion-evoked image stimulus. After a 2000 ms fixation phase, in the evaluation phase participants were presented with the Self-Assessment Manikin (SAM; Bradley & Lang, 1994), and were asked to select “the one that best corresponded to their current state” using a 9-point scale from 1 to 9 for both arousal and valence. First, the participants evaluated “arousal” by pressing the corresponding keyboard buttons from 1 to 9 and then they moved to the “valence” screen. After both evaluations were completed, participants moved on to the next trial. Twenty-five trials were conducted in each condition. Our experimental design followed the paradigm used in previous studies (Mehmood & Lee, 2015; Frantzidis et al., 2010). The “Without Evaluation” condition was based on the literature (Singh et al., 2019) and involved the manipulation of a simple button-pressing task instead of the emotion evaluation task. Immediately prior to the first “With Evaluation” condition (experimental group Block 2, control group Block 4), a practice trial of the “With Evaluation” condition was conducted for three trials, including teaching. At that time, when the SAM was displayed, participants were instructed to “intuitively choose the doll that most closely resembles your current state”. Participants were allowed to freely blink throughout the experiment.

Under the “Without Evaluation” conditions, the evaluation phase was replaced by a button-pressing phase (Figure 2A) so that a 2 s fixation phase after IAPS stimulation was followed by a button-pressing phase. During the button-pressing phase, participants were asked twice to press a key on the computer corresponding to a number from 1 to 9 displayed on the screen. The “Without Evaluation” condition followed the same sequence as that conducted “With Evaluation” condition, except that the evaluation phase was replaced by the button-pressing phase. The numbers were presented in a pre-determined random order and in the same way for all participants, but in the “Without Evaluation” Condition for the control group in Block 2, the numbers were used that had been answered in the “With Evaluation” Condition in Block 2 for the experimental group, which took place one participants before the control group. A simple button-pressing phase under “Without Evaluation” conditions and number control in Block 2 was set up for the purpose of controlling for keypress myoelectricity and button spatial search that were not relevant to the evaluation between groups.

The 100 selected images were presented to the participants in 25 trials x 4 Blocks in the same order in a pre-randomized permutation. This ensured that the images were always novel to the participants. Table 1 shows the mean and standard deviation of the IAPS ratings and the ratings by the participants for each group of images in each Block. The behavioral index was calculated by averaging the rating values for the 25 images recorded in each Block, which was then used as the rating value for each participant.

Participants were provided information about the number and length of experiments but no information about the conditions or order of conditions prior to the experiment. When a condition was new, they practiced three trials prior to its implementation. That is, just before performing the experiment under “With Evaluation” conditions, participants were presented with the SAM for the first time and instructed to select the one that was most equivalent to their current state. During the practice trials, participants were presented with three IAPS images that were not included in the test blocks. These three images were common to all practice trials. Before the start of each block, EEG measurements were taken for 90 seconds each in the order of open and closed eye rest. Each block lasted approximately 4 minutes. In between each block, there was a 1–5 minute break during which the participants were not required to do anything. Participants were briefed about the experiment for 10-15 minutes before the experiment, and practice trials were conducted. The evaluation phase was not included in the data for analysis.

### 2.5. EEG Preprocessing

EEG signals were acquired using an EEG system with 29 scalp electrodes, using Active dry Ag-AgCL electrodes (Quick-30; Cognionics, San Diego, CA, USA). Electrode positions were arranged according to the international 10-20 system. Data from the left earlobe were measured for later offline reference. Signals from all electrodes were sampled at a rate of 500 Hz.

As a pre-processing of the data, a Finite Impulse Response filter of 1–30Hz was applied to the data of each block. The continuous time series data for each electrode was divided for each trial into epochs starting 1000 ms before the onset of stimulus presentation and 3500 ms after stimulus disappeared (total 4500 ms). At each electrode, a semi-automatic procedure of clean_rawdata, an eeglab plug-in, was used to remove noisy electrodes or electrodes that did not provide adequate measurements, and then line noise removal was performed using pop_cleanline, also from eeglab. Noisy epochs were then removed semi-automatically. To be precise regarding epoch rejection, a probability threshold of −500 μV to 500 μV amplitude threshold, 6 standard deviations for a single channel, and 2 standard deviations for all channels was applied. Probability thresholding was performed many times until no data outside the threshold was detected. Independent component analysis was performed on the epochs data to identify and remove artifacts related to eye movements and electrode noise based on component topography and power spectra. P8 electrode data from 10 participants were not recorded due to a hardware misconfiguration, but we treated them in the same manner as noisy or poorly measured electrodes (removed in preprocessing). The data from 9 participants were removed due to severe baseline noise and blink-related noise in almost all trials. The final analysis consisted of clean epoch data from 31 participants (experimental group: 11 male/ 3 female participants, 21.86 ± 4.22 years; control group: 9 male/ 8 female participants, 21.06 ± 0.97 years). The mean number of trials per condition remaining after the artifact correction procedure was 18.27 ± 2.87 (mean ± SD).

All data preprocessing and analysis were performed using custom-written MATLAB scripts and EEGLAB, an open-source toolbox for EEG data analysis (Delorme & Makeig, 2004).

### 2.6. FAA/ERSP Analysis

FAAs were calculated based on Event-Related Spectral Perturbations (ERSPs), which are the results of time-frequency analysis. ERSPs were calculated every 17.5 Hz for each EEG epoch of F3 and F4 electrodes based on eeglab’s pop_newtimef and baseline correction was made by subtracting the mean value of −1000 ms to 0 ms of stimulus onset, the data were converted to left-right differences by log(F4)-log(F3). Frequency power was normalized by this calculation (Sikka et al., 2019). The waveforms for each Block (Block2 and Block3) in the experimental and control groups were visualized by calculating the average amplitude of 8-13 Hz per time. We applied the most common formula used in previous studies for FAA (Vincent et al., 2021).

We examined the effects of evaluations on the FAA by conducting Welch’s t-test on FAA in Block 2 (With/Without Evaluation) for the experimental and control groups. In addition, Welch’s t-test was conducted on the FAA in Block 3 (Without/Without Evaluation) for the experimental and control groups to examine the after-effect of the evaluation on the FAA. The effect size of Cohen’s d value was calculated to show the difference between the groups.

### 2.7. ERP Analysis

ERPs were calculated at the EEG epoch of the Pz electrode and corrected using a reference to the pre-stimulus baseline of −100 ms to 0 ms. The time window for identifying the P300 peak was set between 300 and 500 ms according to a previous ERP review paper (Olofsson et al., 2008) and an article calculating ERPs for similar stimuli (Cano et al., 2009). The average peak amplitude for each participant was obtained by averaging the data over the time interval.

## 3. Results

### 3.1. Induction of negative emotion by IAPS

Before starting the EEG analysis, we confirmed that the stimulus induced the negative arousal emotion as expected evenly in each group and each block. In Block 4, all participant ratings were significantly more negative than the neutral value of 5 (Valence: t (24) =12.29, p<0.001, d=3.48, Arousal: t (24) =-0.42, p=0.68, d=-0.12).

Welch’s t-tests were conducted on each participant’s ratings of arousal and emotional valence for Block 4 in the experimental and control groups and confirmed that there were no group differences in the elicited emotions (valence: t (30) =-1.02, p=0.31, d=-0.36, arousal: t (30) =-0.34, p= 0.73, d=-0.12). We also performed the same test between Block 2 and 4 in the experimental group and confirmed that there was no significant difference here (valence: t (26) =0.26, p= 0.79, d= 0.10, arousal: t (26) =0.07, p=0.94, d=0.03) (Figure 3).

**Figure. 3.**
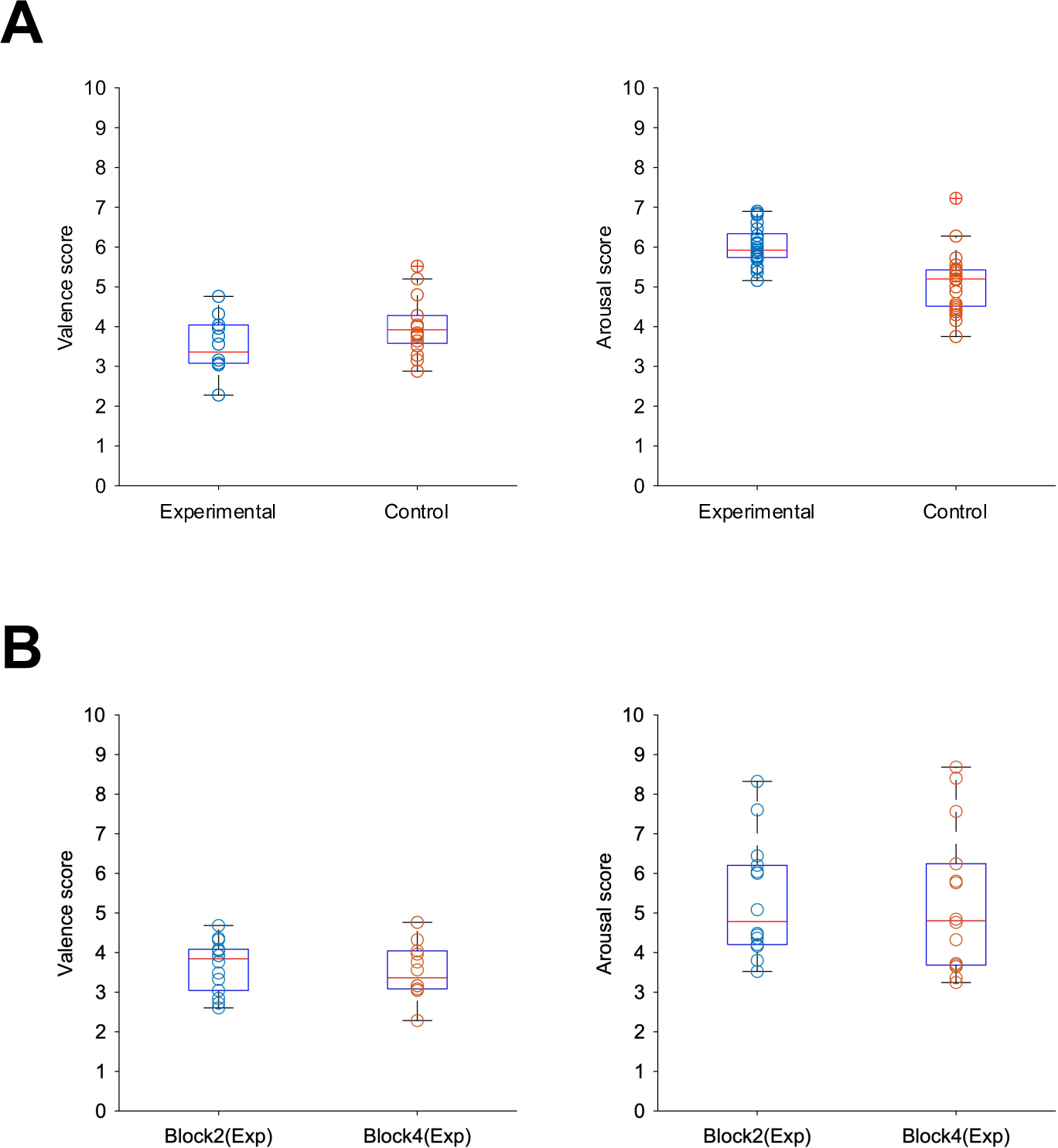
Comparison of participant evaluation values in the experimental and control groups, Block 2 and Block 4. Left: Emotional values. Right: Arousal level. **(A)** Box plots of the average emotional valence and arousal level per participant in Block 4 for the experimental and control groups. No significant differences were detected in emotional valence or arousal (p>0.05). **(B)** Box plots of mean emotional valence and arousal in Block 2 and Block 4 for each participant in the experimental group. No significant differences were detected in emotional valence or arousal (p>0.05).

### 3.2. FAA/ERSP

The FAA results are shown in Figure 4. Figure 4A plots the average time course of FAA across participants for the experimental group (blue) and control group (orange) in the Block2 and Block3, respectively, with the 95% confidence intervals. On the horizontal axis, time 0 is the stimulus onset. Our main hypothesis concerning differences in FAA is that differences are due to the presence or absence of emotional evaluation and experience. Since only negative stimuli are presented in this experiment, if the more the emotion was induced by the evaluation, the smaller the FAA value would be. Welch’s t-test in the mean of FAA between the groups during the 0–1000 ms stimulus interval revealed that FAA was significantly smaller under “With Evaluation” conditions of experimental group than under “Without Evaluation” conditions of control group (t (23.75) = −2.55, p = 0.02, d = −0.95). No significant difference was identified for the after-effect of the evaluation (t (29.00) = −0.19, p=0.85, d = −0.07) (Figure 4B).

**Figure. 4.**
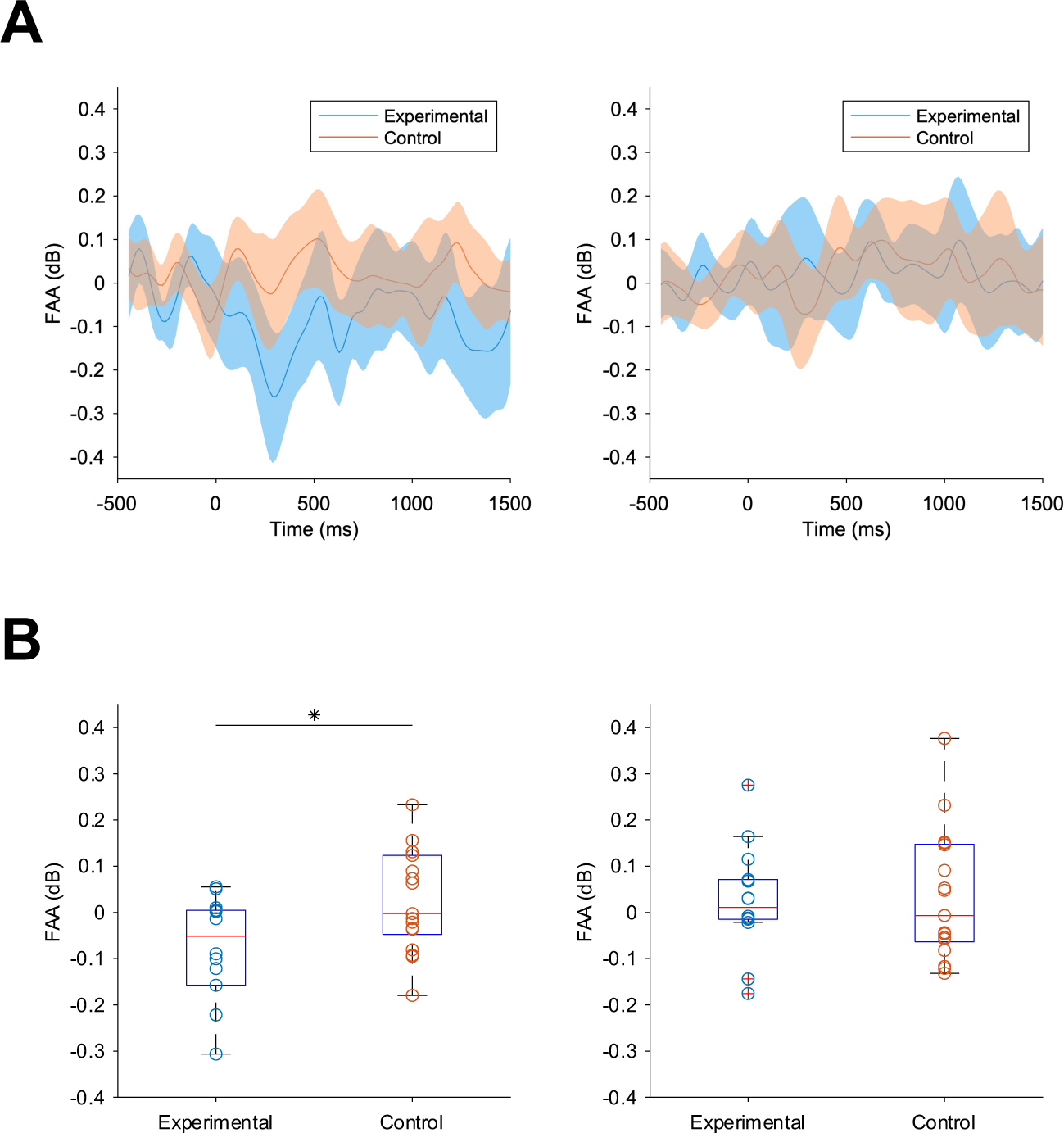
Comparison of FAA between experimental and control groups. Left: evaluation effect (Block 2), right: after-evaluation effect (Block 3). **(A)** Line plots of FAA time variation were calculated based on ERSP analysis for the experimental and control groups in Block 2 and Block 3, respectively, with the 95% confidence intervals. **(B)**: Box plots of FAA for each group in Block 2 and Block 3 at the stimulus interval of 0–1000 ms. There was a significant difference in FAA between the experimental and control groups in Block 2 (p<0.05).

### 3.3. ERP

The ERP waveforms in Pz, in which 0 is stimulus onset, are shown in Figure 5. Figure 5A shows the average waveforms across participants in the experimental and control groups for Block2 and Block3. Comparing the amplitude of P300 in Blocks 2 and 3 between the groups, no significant differences were identified in either Block (t (27.94) = −1.58, p = 0.12, d = −0.57) (t (28.20) = −0.48, p = 0.63, d = −0.17).

**Figure. 5.**
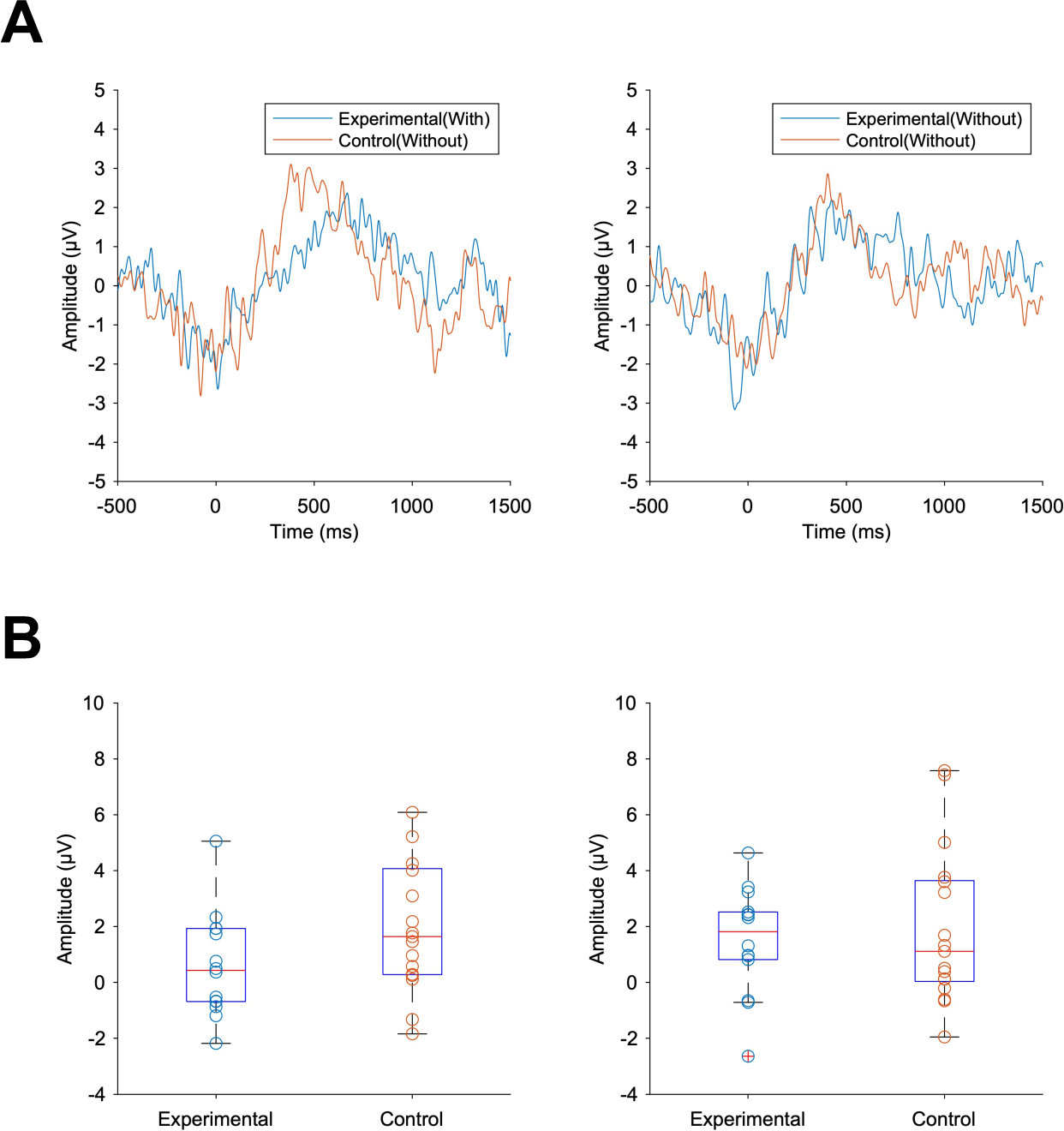
Comparison of ERP waveforms and P300 amplitude and latency between experimental and control groups. Left: evaluation effect, right: after-evaluation effect. **(A)** The plot of time variation of amplitude in Block 2 and Block 3 for each group was calculated based on ERP analysis. **(B)** Box plot of P300 shows averaged amplitude during 300∼500 ms after the stimulus in Block 2, Block 3. There were no significant differences (p>0.05).

## 4. Discussion

The purpose of this study was to determine whether the subjective evaluation of one’s emotional state has an effect on EEG measured FAA during emotion elicitation by visual stimulus or an after-effect on it. In the experiment, two conditions were applied: a condition in which participants were required to make an evaluation of their emotions after stimulus presentation, and a condition in which no evaluation was required. The results of the experiment confirmed that FAA was significantly lower under “With Evaluation” conditions. No after-effects of FAA were identified.

Our main hypothesis was that the FAA elicited greater modulation under “With Evaluation” conditions than under “Without Evaluation” conditions because participants experienced the emotion more intensely as they focused their attention on their own emotions. We also hypothesized that after-effect of evaluation existed. The results supported the hypothesis that the FAA is amplified by the evaluation, but the effect does not continue after the session. In addition, an analysis was conducted in this experiment to see if modulation was confirmed for P300 as in the study by Singh et al. (2019); however, no amplification of P300 was observed. Therefore, we postulate that a modulation of attention occurred in our study than in Singh et al. (2019).

fMRI studies have shown that brain regions associated with emotion are fundamentally altered by observing and evaluating the emotion itself. Lee et al. (2012) summarized fMRI studies showing brain regions associated with emotion evaluation and reported that the brain regions associated with emotion evaluation are the amygdala, lateral prefrontal cortex and dorsomedial prefrontal cortex. The amygdala and prefrontal cortex are brain regions that have been reported to be related to emotion (Davidson et al., 1999, Canli et al., 1998), and thus the results of the study by Lee et al. (2012) suggest that emotion evaluation may alter emotion itself. And Hutcherson et al. (2005) reported that the activation of the anterior cingulate and insula were increased when the participants were asked to continuously evaluate their own emotions while watching an emotion-evoking video, compared to when the participants were asked to simply watch an emotion-evoking video.

While fMRI research has investigated the effects of emotion evaluation, this theme has been overlooked by the increasing number of EEG-based emotion classification studies over recent years. Interestingly, Brouwer et al. (2015) emphasize the need to conduct emotion evaluation during the presentation of emotional stimuli to confirm that the expected emotion is being properly recalled. Our results suggests that there is a critical difference in EEG indices between measurement in an experiment where evaluation is done and measurement in daily life where evaluation is not done. Muta & Koshikawa (2015) reported that answering The Positive and Negative Affect Schedule (Watson et al., 1988) questionnaire in advance influenced the reaction time of a subsequent emotional word discrimination task. They argue that evaluation increases attention to internal states and enhances self-focused attention. Our evaluation task may similarly enhance self-focused attention, and lead to a state of enhanced emotional experience (Scheier & Carver, 1977), resulting in amplified FAA.

In this study, we showed that subjective evaluations of emotions elicited a significant decrease in FAA in the negative stimulus condition; since FAA is a measure that changes in relation to emotional valence, this result indicates that emotions were boosted by the evaluations. Many studies examining the relationship between brain activity and emotion are targeting evaluations for the emotion that the stimulus possesses (Gorno-Tempini et al., 2001, Williams et al., 2006). In contrast, our current study focuses on the evaluation of one’s own emotions during the presentation of emotional stimulus, from the perspective of applying estimation techniques to emotions experienced in everyday environments. Lee et al. (2012) reported that there are differences in the relevant brain regions when evaluating an emotion presented in the stimulus or self-emotion evoked by the stimulus. However, our results regarding the reduction of FAA with emotion evaluation imply a relative right frontal activation with emotion evaluation, which is consistent with those of Gorno-Tempini et al. (2001), who reported right frontal activity during a task to evaluate facial expressions (emotions) in a facial image.

Figure 4A shows that the difference in FAA was more pronounced around 300 ms. after the onset of stimulus. This relatively early appearance of difference in FFA suggests that evaluation affect primary emotional processing including preparation/readiness toward the stimulus, rather than slow, higher-order processing. This quick and transient difference may not be identified by fMRI studies with lower temporal resolution. Regarding the after effect, no difference was observed, which was different from Muta & Koshikawa (2015) reporting that answering The Positive and Negative Affect Schedule (Watson et al., 1988) in advance influenced the reaction time of a subsequent emotional word discrimination task. This result may suggest either that was a difference in reaction time but not in brain activity, or that the after effect did not occur in the first place in the present task.

The results for ERPs showed no significant modulation with respect to P300 amplitude. There are two possible explanations for the difference in P300 modulation between our results and those of Singh et al. (2019). One explanation is that the differences are due to whether participants evaluated the emotion presented in the stimuli or evaluated their own emotion elicited by the stimuli. Singh et al., (2019) confirmed that evaluating emotion presented in the stimuli elicited attention to the stimuli and thus an increase in ERP amplitude appears (Singh et al., 2019). ERP amplitudes for emotional stimuli are reported to decrease when attention is directed to another object or space (Norberg & Wiens, 2013). It is possible that in the present experiment, the condition in which participants were asked to evaluate their own emotional state, attention to the stimulus was reduced as attention was directed to their own state, and the amplification of P300 did not occur. Another explanation is that the emotional stimuli in this experiment were uniformly negative high arousal, and this may have caused habituation or anticipation of the stimuli. In fact, the mean evaluations of arousal for all participants for each image did not differ significantly from the neutral evaluation of 5. As shown in the classical oddball task, ERPs show greater amplitude during deviant stimuli than during familiar and predictable stimuli (Squires et al., 1975). Therefore, we speculate that the ERP amplitudes for the stimuli in this experiment were small to begin with.

There are limitations to this study. First, only negative stimuli were used in this experiment owing to the limited time available to keep wearing the EEG instrument. Since positive and negative emotions may not be symmetrical, it is necessary to separately confirm whether similar changes are observed for positive stimuli in the future study. In addition, a simple button-pressing task was performed as a condition without evaluation in this experiment. This is because we interpret that attention to the internal state is the part of the act of evaluation. However, if we want to confirm the purer effect of emotion evaluation, we need to have a control condition with the operation of paying attention to the internal state without evaluating the self-emotion.

## 5. Conclusion

In this study, we found that FAA was significantly lower in the emotion subjective evaluation condition during negative emotion evoking stimulus presentation. This result highlights the possibility that subjective evaluation of their own emotion may have an immediate effect of emotion amplification. The results suggest that evaluation may skew EEG during emotion recognition and call for caution in considering study methodology in emotion recognition research.

## Declarations of interest

None.

## Funding

This work was supported by JSPS Grant-in-Aid for Scientific Research(A) Grant Number 19H00635.

## Data availability statement

The datasets generated and analyzed during the current study are available from the corresponding author on reasonable request.

## Credit Author Statement

**Masato Ito**: Conceptualization, Data curation, Formal Analysis, Investigation, Methodology, Software, Visualization, Writing – original draft, Writing – review & editing. **Toru Takahashi:** Conceptualization, Methodology, Writing - review & editing. **Yuto Kurihara:** Methodology, Writing - review & editing. **Rieko Osu:** Conceptualization, Funding acquisition, Methodology, Supervision, Validation, Writing - review & editing.

## Notes

### Competing Interest Statement

The authors have declared no competing interest.

